# Microfluidic Tissue Array Platform for Personalized Drug Screening Using Tumor Explants or Biopsies

**DOI:** 10.64898/2026.07.01.735821

**Authors:** A.H.R. Ahmed, H. Shao, L. Colón-Cartega, L. Wang, X. Jiang, F. Pareja, S. Chandarlapaty, S. Wang

## Abstract

Despite major improvements in molecular characterization of breast cancer, current biomarkers still fall short in accurate treatment prediction. Interrogating tumor tissue ex-vivo in its native conformation is a direct strategy for guiding treatment of individual patients but presents a challenge. In this study, we developed a microfluidic tissue array (μFTA) using small biopsy samples (< 1mm3) mimicking physiological flow for consistent exchange of nutrients and waste, retaining the tumor native stroma. Cell/patient-derived breast cancer xenograft tissues were maintained over 2 weeks in the array and their response to therapeutic agents, doxorubicin or neratinib, were interrogated. Drug response in the uFTA showed >2-fold reduction in tumor cell viability which corroborated tumor size shrinkage in mice bearing the same tumor load. EdU/Ki67 assays indicated selective retention of cells with higher proliferative capacity after drug treatment, underscoring in vivo clinical relevance . We have also developed a valved-μFTA to increase throughput and variety of treatment conditions on the same chip. Together, this μFTA can be staged as a powerful, low-cost benchtop theranostic tool for personalized cancer therapeutics compatible with FDA’s New Approach Methods.

## Introduction

Cancer is a multifaceted disease resulting from the expression of accumulated genetic mutations and epigenetic triggers. The heterogenous complexity of a whole tumor microenvironment implies that drug screening platforms for cancers cannot be limited to traditional cell line models using homogenous populations of cells. Cancer treatment itself can be highly multi-modal, and cancer drugs have a low 3.4% success rate of approval from Phase 1 clinical trials [1]. Traditional 2D cell culture platforms do not facilitate understanding the resistance or adaptation to drugs that cells in a 3D environment modulate based on interactions with other cellular and non-cellular stromal partners [1]. Cellular components modulate drug sensitivity, especially cancer associated fibroblasts and endothelial cells[2]. Non-cellular components, including alterations in the matrix stiffness, biochemical gradients and interstitial flow/pressure properties, also modulate tumor growth and maintenance. [3–6] A holistic reconstruction of these structural and functional environmental cues are crucial to obtaining more faithful trajectories of drug treatment regimen. The ease of processing, handling, and scaling up of 2D cultures cannot override the prognostic reliability from a fully perfused 3D system.

### 3D culture models to address 2D limitations

Some limitations of traditional 2D cultures are being addressed with emerging 3D tumor models that recapitulate or do not disrupt physiological tumors. Recapitulation typically involves formulation of the optimal extracellular matrix (ECM) within which isolated tumor and supporting cells are reconstituted [4,7]. Currently recognized models include patient derived xenograft (PDX) and patient derived (cancer) organoids (PDO/PDCO), which harness the 3D format of tumor tissue microarchitecture. PDX are generated in animals from patient tumorigenic cells [8–10], while PDOs use patient-derived cancer/stem cell population to generate a heterogeneous tumor with a mix of aptly differentiated cell types[9,11–15]. Chemosensitivity to standard therapeutic drugs can bear a good correlation to patients’ clinical outcome from whose tumors the PDX or PDCOs are generated, although a significant change in progressions-free survival (PFS) or overall survival (OS) is not predicted by these models [16]. The newer approach of tumor spheroids enjoys a prolific utility due to ease of generation compared to a complete PDX/PDCO model [13,15,17,18], as spheroids can be generated from highly proliferating cancer cells without using special stem-cell differentiation media or waiting for differentiation into a fully heterogeneously populated tumor organoid over long time [3,13,16–18].

On the other hand, approaches relying on maintenance of the environment utilize the whole tumor in its original excised form, cultured as explants after mechanical reduction to a working size, and are supplemented with appropriately optimized media formulations - a prevalent procedure exercised in *ex vivo* organotypic cultures (explant or micro-dissected types) [19–23]. Organotypic cultures rely on directly culturing tumor explants ex vivo, and direct histoculture of patient derived explants (PDEs) - or their animal analogs in preliminary trials - have been employed since the 1960s [24–28].

### Microfluidics-based 3D Culture Systems

Emerging paradigms of patient-specific drug testing using microfluidics have also been in the forefront over the last decade, including microfluidics-based culture (better known as organs-on-chips) and micro-organospheres (MOS). One of the more famous organ-on-a-chip systems is the lung-on-a-chip [29]. MOS systems utilize the fluidic manipulatory capability of microfluidic systems to combine microfluidics and PDOs/PDCOs, generating mini-ecosystems of organoids within sorted droplets. As with PDOs/PDCOs or PDEs, early passages contain stromal cells, and allow for various immune checkpoint assays as well.[30]

### 3D models are inadequate for personalized medicine

All the current standard models - 2D culture, spheroids, PDOs/PDCOs, PDEs, Organ-on-chip, and MOS systems - pose their own burden in modeling clinical outcomes for patients. For these 3D models that are reconstructed using cell lines or isolated primary cells with hydrogel, their homogeneously bioartificial design lacks the complexity of native tumor microenvironment (TME) and thus is not sufficient for the evaluation of drug responses and is not recognized by insurance companies [31]. In the case of PDX models, tumors in animals do not always establish successfully, being as low as 21-35% success rate for breast cancer [32], while highly metastatic forms (e.g., the breast cancer metastasis into the brain) from certain patients could be engrafted with a 100% uptake rate [8]. Although over 90% tumor engraftment success has been reported for other cancer type/mouse combinations, including prostate, non-small cell lung, and head and neck cancers in NSG, NOG, or NOD/SCID mice, the typical success rate has a wide variance from 21-100%, and is influenced by individual patients, tumor acquisition method, host strain, implantation site, sample volume, carrier matrix, and hormonal supplementation [8,32]. Successful establishment in itself can be a lengthy process. For PDX, throughput by design is low due to in-vivo growth time in mice. Broad variation in successful establishment accompanies PDO generation as well, varying from 15% to 100% across cancer types [33], with the tumorigenic content of the originating sample being a strong determinant in the process (e.g. for breast cancer tissue there is >80% success rate vs ∼20% for healthy tissue[14]). Multi-omics screening can provide some characteristic drug regimen, but is not always fully predictive[34]. In instances where multi-omics screening does not provide a clear selection, off-label use of agents are routinely considered and may be covered by insurance if the drug is listed in an approved compendium[35,36]. All these issues necessitate the development of a new model that : (1) recapitulates tumor tissue structure and heterogeneity, (2) permits rapid but clinically relevant mid-to-high throughput assay systems to evaluate drug or combination treatment efficacy, (3) deliver the screening results in weeks instead of months, and (4) has a lower cost than PDX/PDO models

### Shift from generative 3D models to direct tissue interrogation

Several ex-vivo methods using intact tumor explants alleviate the aforementioned shortcomings of 3D platforms . Ex-vivo platforms include using resized tissue blocks, slices, or cored biopsies in: (1) standard cell/tissue cultureware with customized media formulations[37], (2) standard cultureware placed on moving platforms for flow[4,19,38], (3) microfluidic systems for tissue trapping[20,39], (4) microfluidic systems with perfusion along a tissue edge[20,40], (4) macrofluidic/microfluidic systems of modified or custom cultureware platforms with mechano-electrical systems promoting perfusion through the tissue pieces[41–43]. These systems can include scaffolds to support the microtissues, using niche geometries of processed tumors in their systems. The geometries vary from blocks or discs a few millimeters across and hundreds of micrometers thick to long, cored out tissues a few centimeters long. These systems have throughput limitations or unreliable fidelity in emulating the controlled perfused nature of natural tissues *in vivo*, showing noticeable loss in viability or functional activity after 24-72 hours of culture [40]. Sustained perfusion is a critical *ex vivo* factor in continued advective delivery of nutrients to deeper regions, and in maintaining interstitial mechanostimulation drivers of tissue growth.

### A new model that conserves tissue architecture and sustains perfusion

Here we develop a platform that allows a mid-throughput culture of whole tumor tissue samples that are microdissected to size (500-1000μm along each dimension). Our patented microfluidic tissue array (μFTA) [44] addresses critical concerns with the previously published *ex vivo* platforms. The μFTA (1) works with 500-1000μm thick *tissue samples*; (2) supplies *continuous* and tunable *perfusion* matching 0.1-10 μm/sec interstitial flow rates, facilitating nutrient, gas, and waste exchange; (3) provides an *automated trapping mechanism* for seeding tumor pieces, (4) permits direct *on-chip viability and proliferation assays for drug treatment*; (5) has a structured design of micro-chambers that enable medium self-conditioning. We highlight direct tumor tissue culture in our μFTA using several xenograft models here, focusing on 2 cell-line derived xenografts (CDX) for initial system validation and 3 patient derived xenografts (PDX) models, with both model sets having a triple negative (HER2-/ER-/PR-) breast cancer (TNBC) or HER2+ types. Doxorubicin (for TNBC) or neratinib (for HER2+) was evaluated for efficacy in the μFTA. Our μFTA was able to maintain explanted PDX samples for two weeks. Overall, the efficacy of drug treatment was also successfully interrogated in our μFTA explant cultures, showing a correlation to *in vivo* treatments in mice.

## Results

### μFTA Device Design, Optimization, and Implementation

*μFTA Manufacture and Utilization.* The μFTA device consisted three layers, top media feeding layer (500-700µm depth), bottom tissue seeding layer (400-600µm depth), and middle layer (40-60µm) with pores (2x2 sets of 200µm-diameter) dispersed to interface flow between top and bottom layers. Our design allows for flowing small tumor pieces of size 0.03-1mm^3^ through the bottom layer and media through the top layer. A trapping mechanism (“catcher”) is integrated into each individual chamber of the bottom layer (Fig. 1.A) with a trapping efficiency of 60-90%. Of several array formats (2x4, 4x4, 4x8) of the µFTA, two 4x4 are shown in Figures 1B-C, with food dye-stained water permeating the channels. The top layer lays out the connectivity for providing perfused flow to the tissues within the bottom layer through the porous middle layer. The flow-driven mechanism for microtissue entrapment shows captured pieces of tissue visible in the catchers of the bottom layer (Fig 1D).

**Figure 1.**
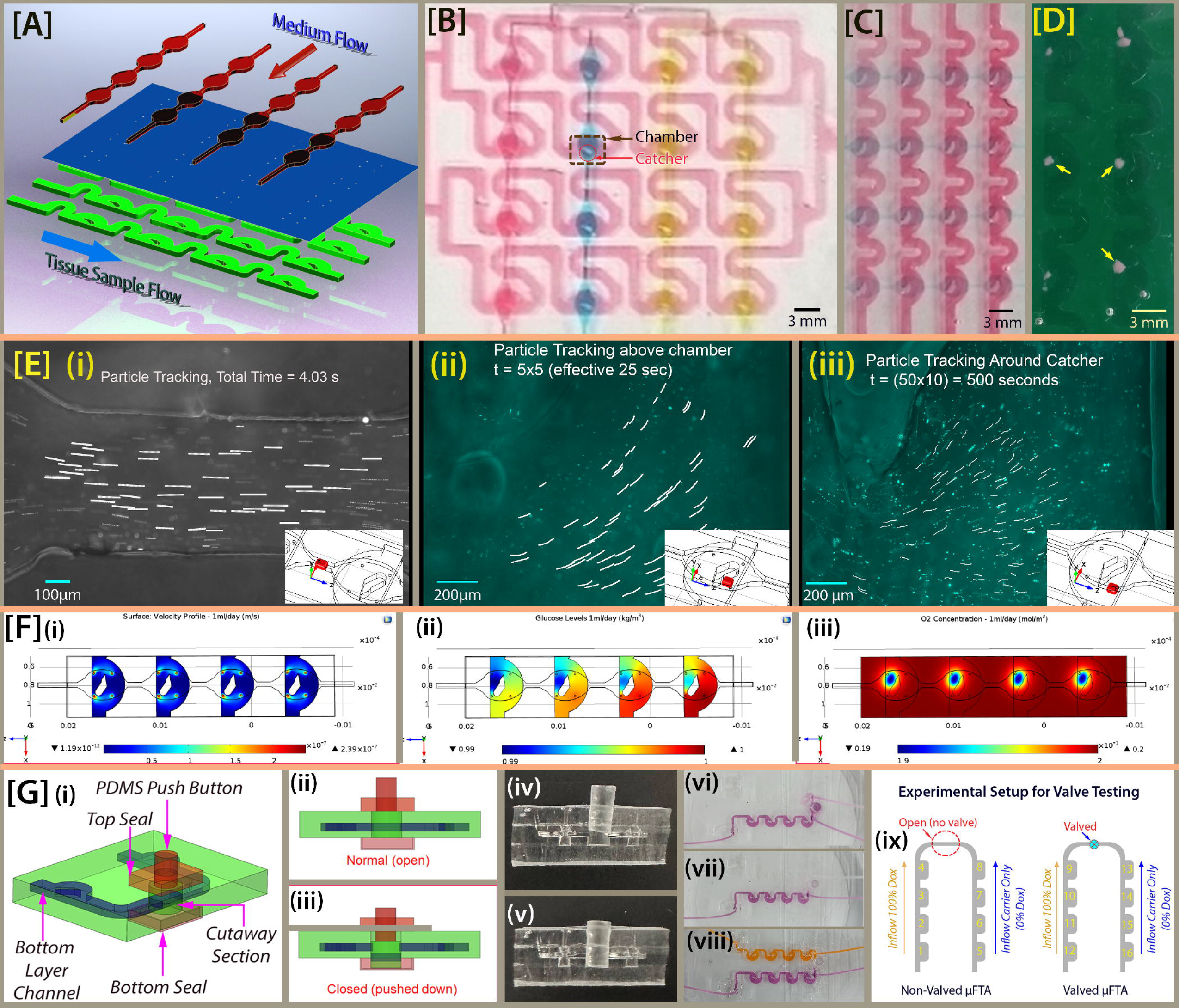
Conceptualization, development, functional testing and performance validation of microfluidic tissue array (μFTA). (A) An exploded conceptual CAD rendering of the layered structure and flow direction in the μFTA, showing the top media flow layer (red), bottom tissue seeding layer (green), and the pore-laden middle layer (blue) interfacing the top and bottom. (B) First generation PDMS prototype of the 4x4 μFTA, with long, meandering, diffusion-limiting channels between chambers. The bottom layer is perfused with red dye, while the top media layer’s individual channels are shown with different dye colors (red, blue, yellow, green, respectively, from left to right). The typical chamber region in the bottom layer and the catcher location are marked. (C) A second-generation μFTA device with reduced footprint. The bottom and top layers are filled with red and blue dyes, respectively. (D) Microdissected tumor samples trapped in the μFTA catchers. (E) (i-iii) Particle tracking velocimetry at different times scales, using fluorescent 2μm latex beads within the perfused media; insets indicate probing locations. Particle traces (i) in the top layer between chambers; (ii) in the top layer above the tissue catcher of the bottom layer; (iii) in the bottom layer at the catcher. (F) CFD results from COMSOL at the bottom layer, showing the (i) velocity profile, (ii) glucose concentration, and (iii) oxygen concentration, under emulated nutrient consumption rates and biotransport factors. (G) Valve prototype and testing, shown first as a (i) CAD rendered model, and an (ii) initially open state and (iii) pushed, closed state. (iv-v) A cutaway cross section of the real, PDMS-push valve in the μFTA showing the (iv) open and (v) closed states. (vi-viii) Valve in action showing cross-talk across the valve using a mix of clear or colored food dyes under flow conditions. (vi) Image with valve open, within 2 minutes of the flow initiating in both channels - top channel running clear water and the bottom running violet food dye in water. (vii) Image of the device in (vi) with closed valve, taken 5 minutes after flow initiation. (viii) Close valve setup with an orange food dye in top channel, and same violet dye in the other channel. Image taken after 5 minutes since flow began. (ix) Schematic representation of a long term valve test setup, to measure detectable cross-talk across the valved chambers over 24 hours

CFD simulation was utilized to optimize the μFTA design and flow rates to simulate *in vivo* interstitial flow around microtissues. Simulation results of 1ml/day flowrate is shown in Figures 1E(i-iii). Figure 1E(i) shows flow velocity around the tissue on the order of 0.1µm/sec (maximum 0.24µm/sec). Higher flow rates (not shown) are obtained proportionally by increased flow rate at the top layer inlet of the µFTA, with 5ml/day and 10ml/day producing maxima ∼1.2 µm/sec and ∼2.4µm/sec respectively around the simulated tissue piece. Figures 1E(ii) and 1E(iii), depicting glucose and oxygen levels around the tissue respectively, indicate the healthy maintenance of both nutrients via continuous perfusion, with glucose dropping by only 1% (1 to 0.99 g/L) and oxygen only by 5% (0.2 to 0.19 mol/m^3^) from the first to the last of the four serially connected tissue culture chambers. Particle image velocimetric (PIV) data based on the 1-10ml/day flow rates validate simulation results as shown in Figures 1F(i-iii), with three regions of interest tracked: (1) the constricted part of the top layers’ media channel (Fig. 1F(i)) tracked for 4.03s, (2) the wider section in the top layer above the catcher for 25 seconds (Fig. 1F(ii)), and (3) directly at the opening of the catcher for 500 seconds (Fig. 1F(iii)). Matching regions of interest (insets) were also probed in the simulation model and measured from the μFTA PIV. The average velocities of the experimentally traced particles were 21.7 (±3.38)µm/s, 2.83 (±0.76)µm/s, and 0.103 (±0.019)µm/s in regions of Figures 1F(i),(ii), and (iii) respectively, agreeing with corresponding simulation results of 22.7 (±3.35)µm/s, 2.73 (±0.32)µm/s, and 0.113(±0.067)µm/s. (Supp. Fig. 2)

The valve mechanism compartmentalizing rows from communicating with each other was implemented as a post-assembly modification µFTA, designed as a one-time push valve button shown schematically in orthographic view (Fig. 1G(i)) alongside operational cross-sectional views for the open (Fig. 1G(ii)) and closed (Fig. 1G(iii)) states. The real valve is shown via a cross-section in the open (Fig. 1G(iv)) and closed (Fig. 1G(v)) states. Leakage across the valve was visually evaluated with clear or dyed water in one channel juxtaposed with dyed water in the adjacent channel across the valve (Fig. 1G(vi-viii)), showing no short- or long-term cross-talk through the closed valve (Fig. 1G(vii,viii)). In the open state, even under steady flow, the violet dye leaks into the clear (water) channel within the first 2 minutes (Fig. 1G(vi)). Once the valve is closed, the leakage is stopped and the two streams do not cross talk, as seen in the well-separated violet/clear (Fig. 1G(vii)) and violet/orange food-dye channels (Fig. 1G(viii)). The channel separation images captured at 5 minutes (Fig. 1G(vii, viii)) stayed stable over 24 hours. Doxorubicin used as an autofluorescent drug to detect cross-talk (Fig 1.G(ix)) in a valved and non-valved system showed no detectable cross-talk across the valve (Chamber# 9) over a 24-hour duration in the valved setup, maintaining doxorubicin below the 0.02% background signal at the 0% Dox input side.

### Growth And Drug Response of Tumor Tissues in µFTA with Functional Assays

#### Cell-Derived Xenograft (CDX) Explant Growth

Microdissected MDA-MB 231-GFP CDX tissue (murine) cultured for a week to assess the μFTA utility shows structural conformation and cell spreading. A sparse population of rounded, stranded cells are seen on Day 2 around the tumor piece [Fig. 2A(i-ii)], but by Days 5-7 [Fig. 2A(iii,v)] a monolayer of spindle-shaped cells forms around the tissue, typical of MDA-MB231 phenotype [insets of Fig. 2A(iv,vi)]. We also observe tissue pieces losing their sharply defined edge and presenting a softer, more rounded edge by Day 5. Within 3 days of doxorubicin treatment, tissues clear out to look translucent under phase contrast microscopy and the monolayer around the tissue is removed, indicating massive cell loss. The cleared regions lack GFP signal, confirming absence of cells [Supp. Fig. 2]. Viability assays of 1 week μFTA culture of wild type MDA-MB 231 (without GFP) revealed an overall 4-fold increase in peripheral viability (p<0.002). For BT474 CDX, there were no statistically strong differences between tissues cultured in the µFTA vs static culture measured through RFP signals or viability assay [Supp. Fig. 4A]. There was also no difference in the tumorigenic cell content between the two conditions, measured as the ratio of RFP to Hoechst stained cells. However, there are strong morphological differences between µFTA [Fig. 2B(ii-iii)] and static [Fig. 2B(iv-v)] conditions, observed in the formation of spontaneous “budding” spheroids on the edge of the tumors within the µFTA. These buds are rich in tumor cells (strong RFP signal), indicating the spontaneous outgrowth of tumorigenic cells not realized in static cultures.

**Figure 2.**
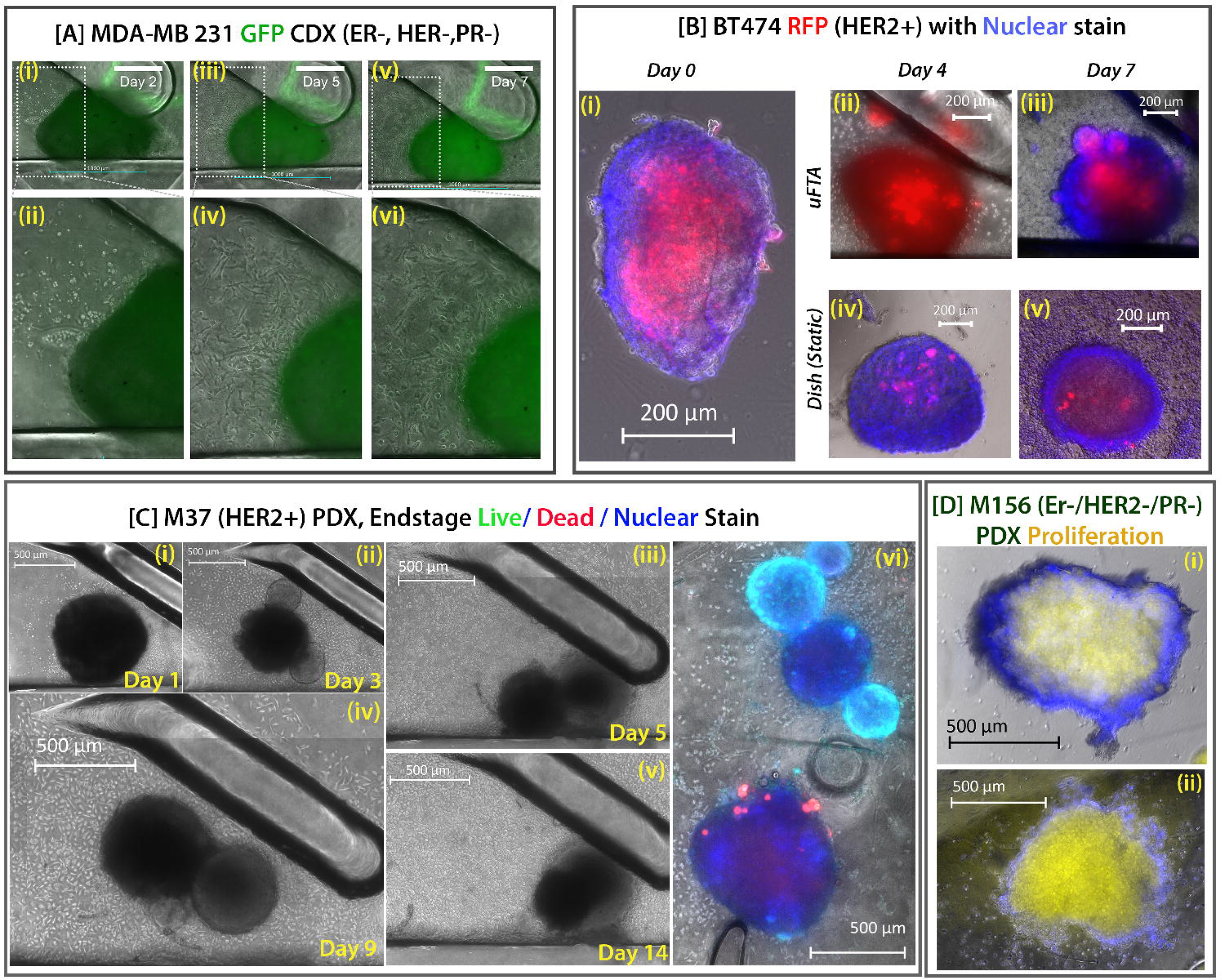
Culture and viability or proliferative characterization of CDX and PDX models in the μFTA. Micrographs of the culture of xenografts generated from the cell lines (A) GFP-expressing MDA-MB231 (triple negative) and (B)RFP-expressing BT474 (HER2+) over the course of a week are shown. (C) A patient-derived xenograft model, M37 (HER2+), was cultured for up to 14 days (i-v), with (vi) end stage viability staining. (D) A triple negative patient derived xenograft, M156, was cultured for a week, and proliferation staining was assayed on (i) day 0 and on (ii) day 7.

#### PDX Explant Growth

A 2-week culture of M37 PDX samples in µFTA show an impetuous growth of spheroid “buds” of highly active and proliferating cells on tissue pieces [Fig. 2C(ii-v)]. In the sample shown, two buds sprouting by Day 3 [Fig. 2C(ii)] shift and merge into a larger, single spheroid by Day 5 [Fig. 2C(iii)], and by Day 14, this super-bud engulfs the originating tissue [Fig. 2C(v)]. In Fig. 2C(vi), the viability staining from the budding piece on top to a non-budding one at the bottom shows a vivid Calcein-AM signal on the buds themselves compared to the rest of the tissues around. M37 samples cultured under static conditions, on a culture plate or PDMS surface, show highly suppressed viability by Day 4 compared to Day 0 (p<0.0001, Supp. Fig. 5A). The µFTA maintains a comparable viability to Day 0, with the peripheral region indicating recovery and increased viability compared to the core (p<0.01). Doxorubicin treatment had no noticeable effect on the viability of M37 [Supp. Fig. 5B], and no negative effect on proliferation (EdU). In M156 PDX samples [Fig. 2D], a 7-day µFTA culture showed two-fold higher proliferative cell content compared to the initial tumor piece [Fig. 2D(ii)], especially closer to the periphery region (N = 9, p<0.01, Supp. Fig. 4(c)). Tissue core viability showed a modest but non-significant improvement.

#### PDX (M271) Explant and in vivo Analogue Drug Responses

On-chip M271 viability and proliferation micrographs are depicted in Figures 3A and 3B, respectively. Following a 72 hour culture period, a 96-hour 1X neratinib (180nM) treatment has a mild, non-significant effect on tumor viability compared to the vehicle [Fig.3A(iii) vs (i)]; however, a 10X dose (1800nM) causes significant cell death within the tumor mass [p<0.001, N=16, Fig. 3A(iv)] seen as large, dead regions marked via Ethidium Homodimer. Data is summarized in Fig. 3D(i). No significant difference in proliferation is seen between the drug vs vehicle samples in the µFTA [Fig. 3D(ii)]. Parallel experiments done *in vivo* in mice indicate similar trends, with all tumor volumes shrinking to <100 mm^3^ for initially 130-200mm^3^ tumors by day 30 of neratinib administration. All non-treated mice retain 125mm^3^-215mm^3^ tumors at the end of 30 days. Notably, the effect of neratinib on treated mice presented after Day 18 [Day 25 in Fig.3D(ii)] of treatment, compared to only 96 hours in the µFTA. Western blot results of excised tumors from the mice show absence of HER2+ expression in neratinib-treated mice. [Total HER2, Fig.3C].

**Figure 3.**
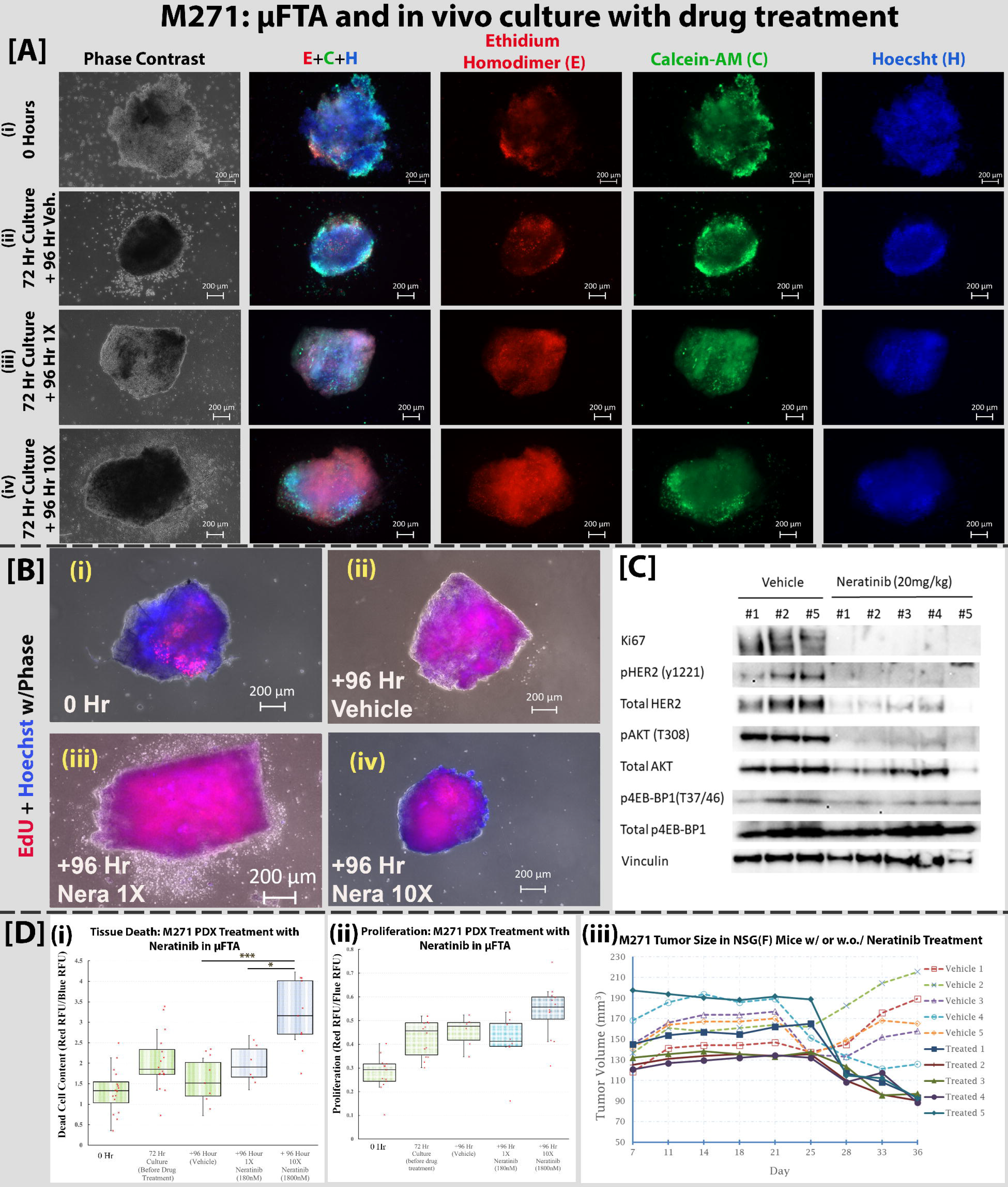
Drug screening of cultured PDX model M271 (HER2+) in the μFTA. M271-derived xenografts were cultured for a period of 3 days (72 hours) followed by 4 days (+96 hours) of treatment with vehicle, 1X or 10X Neratinib concentration. (A) Viability staining of the (i) initial tumor state, (ii) vehicle, (iii) 1X neratinib, and (iv) 10X neratinib are shown, with the corresponding phase contrast micrograph in the first column. (B) Proliferation assay via EdU staining kit is (i) initial tumor state, (ii) vehicle, (iii) 1X neratinib, and (iv) 10X neratinib as well, corresponding to the same time points as the viability stain in (A). (C) Western blot assay from the in-vivo arm of the neratinib treated/non-treated mice indicating the high presence of Ki67 and HER2 in mice receiving no treatment (vehicle) compared to neratinib-treated mice. (D) Fluorescence assay analysis via in-house image processing algorithms, measuring the (i) noticeably increased cell death with increased drug dose (10X), and (ii) no statistically significant shift in proliferation over the assayed time points (0 hours, 72 hours, +96 hours of vehicle/drug treatment). (iii) Measurement of the in vivo tumor xenograft size in mice. [*: p<5%, ***:p<1%]

**Figure 4.**
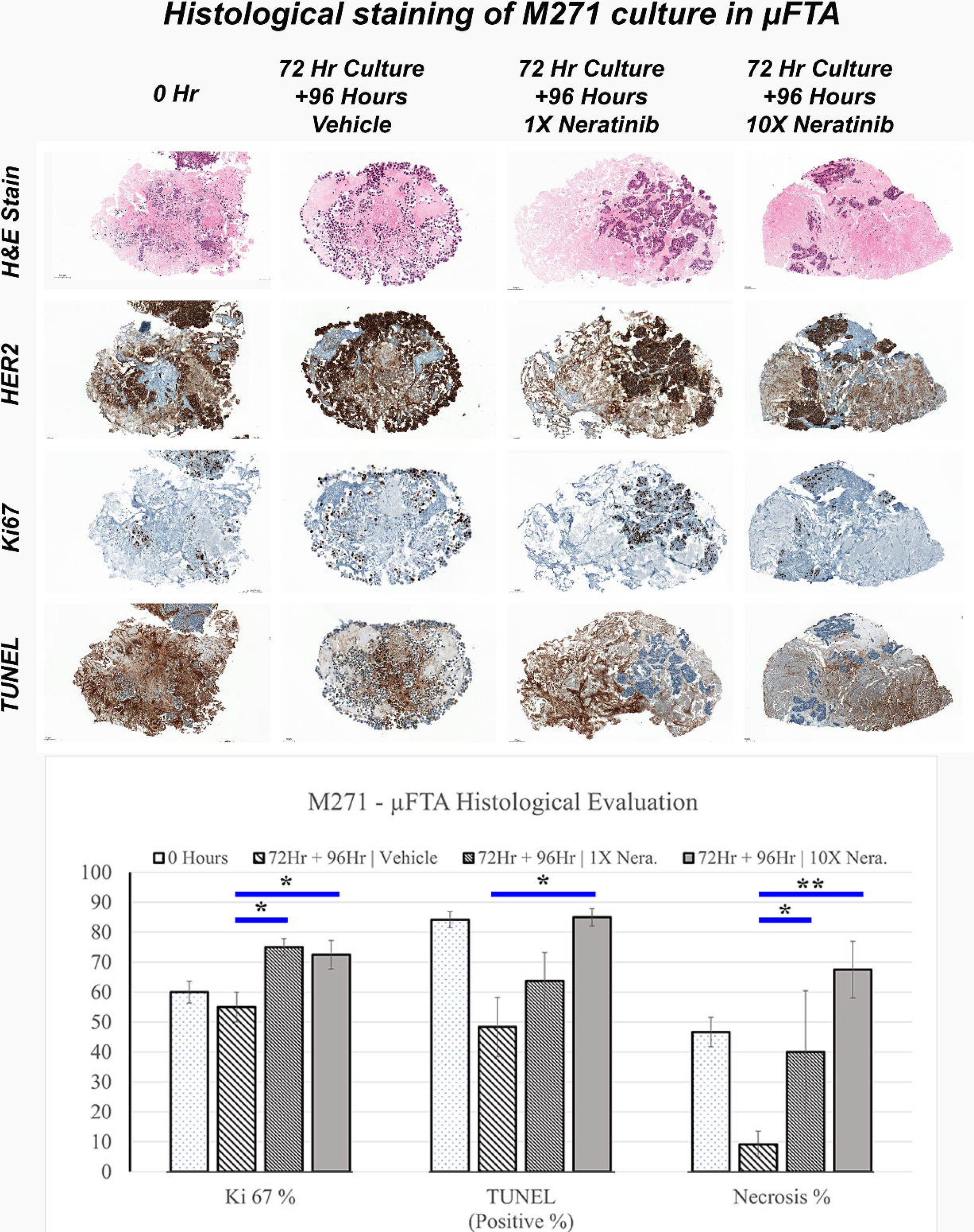
Immunohistochemical evaluation of M271 microdissected tissue cultured and treated in the μFTA. **(A)** H&E, HER2, Ki67, and TUNEL staining of representative samples of each condition going from left to right columns are shown, with the initial state of the tumor (Day 0), followed by a 3 day culture with 4 day vehicle treatment (72Hr + 96 hours Vehicle), 3 day culture with 4 day treatment of 1X dose of Neratinib (72Hr + 96 hours 1X Neratinib), and 3 day culture with 4 day treatment of 10X dose of Neratinib (72Hr + 96 hours 10X Neratinib). H&E stains indicate loss of cellularity in drug treated samples. Strong HER2 expression is visible across cells present within the tumor. Ki67 and TUNEL data are summarized in **(B)**, wherein the change in Ki67 (proliferation) is mild between the drug-treated and vehicle samples, but both TUNEL (cell apoptosis) and necrosis area present a consistent increase with increasing drug dose. (N = 20; n(Day 0) = 6, n(Vehicle) = 6, n(1X) = 4, n(10X) = 4; *: p<5%, **:p<1%,).

#### M271: Histology/Immunostaining

Histological staining of µFTA-cultured tumor samples corroborate in-device viability/proliferation assay results [Fig 4B]. Ki67 indicating cell proliferation shows a significant (P<0.05) increase by about 30-45% (∼32% and 36% between average values) between untreated (vehicle) vs drug-treated (1X or 10X) conditions for the same 7-day period [Fig 4.B, Ki67%]. TUNEL staining indicates a cellular death rate accentuated by drug treatment, (P<0.05) ramping up TUNEL expression by 75% under 10X Neratinib, while a statistically insignificant change of ∼32% is seen in the 1X treated samples[Fig 4B, TUNEL%]. Severely elevated necrosis levels occur in Neratinib treated samples compared to low necrosis (∼9%) in untreated tissue at Day 7: 1X Neratinib shows a necrosis level of ∼40% (P<0.05), and 10X Neratinib at ∼68% (P<0.01) [Fig 4.B, Necrosis%].

## Discussion and Conclusion

The μFTA chip was designed to work with microdissected tumors of size <1 mm^3^, with a minimum of 0.5mm along one dimension. We used optically clear, gas-permeable, biocompatible PDMS (polydimethylsiloxane) commonly employed in BioMEMS to facilitate gas exchange and imaging. Our μFTA design includes catchers that entrap tissue samples automatically as they are carried via flow. Design optimization is done through computational models and particle velocimetry verification to mimic the physiological flow regime around the tumor tissue (0.1-1μm/sec), to theoretically keep the tumors in their most native mechano-stimulatory state. Separating seeding and feeding layers allow independent tuning of each, with a defined porous zone connecting them for nutrient and waste exchange. The tunability of the flow profile is one of the major strengths of the μFTA compared to contemporary organotypic tissue culture platforms. [4,20,48]

Development of cancer therapies utilize *ex vivo* modalities, including tissue culture plates, transwells, microfluidic networks, or other custom-made culture platforms, available in 2D or 3D formats. 2D cultures are easy to handle but the results can severely detract from realistic outcomes, often contradictory to their 3D counterparts[1,5,7,49]. 3D cultures entail many formats - from the simplest cell-in-gel suspensions or spheroids to meticulously generated organoids, animal xenograft models, or complex pre-designed microfluidic architectures. While both patient-derived xenografts (PDX) or organoids (PDOs) can serve as concordant analogs to indicate treatment prognosis, model development can be arduously long, averaging up to 3-12 weeks for PDOs or PDXs inclusive of drug testing [9,12,14,16,32]. Other detrimental factors impacting these models are the digestion processes that remove the originating extracellular matrix (ECM), and reconstitute cells in foreign matrix (e.g. Matrigel) with custom growth factors (in PDOs), or, replaced by animal (non-human) ECM in PDX. Variable success rates in tumor establishment of PDX preclude high-yield capacity for personalized medicine. Regardless, PDX and PDO formats current *ex vivo* gold standards for the closest *in vivo* mimics.

Direct testing of primary tumor tissues without disseminating the integral architecture of the original tissue is a vital step up from PDX/PDO models. Successful culture and therapeutic interrogation of primary tumor tissues should (1) reduce time and failure rate for tumor generation as a sustained culture; (2) preserve, at least short term, all cellular and extracellular components (including immune cell populations); (3) indicate functional drug response directly from a patient tumor to adjudicate well-tailored personalized therapeutics. Several works incorporating whole tissue using custom-made devices or defined culture protocols lack well-characterized mechanical (flow) and nutrient exchange parameters expected *in vivo*. Our work here addresses these shortcomings by engineering a μFTA which provides (1) physiologically matched continuous flow (0.1-1μm/sec) for mechanostimulation, (2) sustained delivery of fresh nutrients or therapeutic agents and removal of waste, (3) ease of automated flow-guided seeding of microdissected tissue pieces into an array for technical replicates, and (4) a culture unit structured for media self-conditioning. Interstitial flow-derived mechanotransduction is a crucial component for healthy or tumor tissue morphogenesis and functional maintenance [50–52], and drug effects rely on proper advective drivers via interstitial flow [53]. The viability and physiological replicability of the *ex vivo* microdissected tissue culture is enhanced by enabling these crucial flow and self-conditioning factors. The μFTA was able to sustain whole tissue from various tumor types over up to 2 weeks, maintaining the primary tumor’s biomarker expression and permitting concordant drug treatment results.

There is a strong incentive for cancer treatment screening technologies that significantly improve quality of life for patients by selecting an efficient treatment regimen conducive to successful remission with minimal recurrence risk tailored to the individual. Current US insurance policies lack support for whole tissue *ex vivo* testing as cost beneficial for personalized medicine coverage, while accepting the potential value of these platforms from several retrospective studies that fall short of tipping the scale beyond existing contemporary methods. Outside of the US, explant culture platforms are accepted by health insurance to be profound screening tools for personalized medicine, one such example in Japan where collagen droplet screening (spheroid cultures, or PDCOs) and histological drug resistance assays (HDRAs using whole explant) has been supported by insurance since 2012[16]. Through future validation and systemic integrations, we anticipate our μFTA to be an everyday diagnostic tool alongside standard assay and sequencing measures to enhance the tailoring of much-needed treatments across individual patients. For validation, more patient samples comparable to their clinical outcomes are necessary. For systemic integrations, the current μFTA will be integrated with our work on a recirculating chip module [30], such that immunological effects can be inserted onto the interrogated biopsy samples. This will add another dimension towards a complete picture of physiological response that entails immunological interactions and biologics, especially involving immune checkpoint interactions.

## Methods

### Media/Drug Formulation

Unless otherwise stated, for all culture conditions, Fetal Bovine Serum, Premium (FBS) was from Atlanta Biologicals, USA; Penicillin-Streptomycin (P/S), Phosphate buffered saline (PBS), high glucose Dulbecco’s Modified Eagle Media (DMEM-HG) from Gibco/Thermo-Fisher Scientific, USA; Doxorubicin Hydrochloride (Dox) and Hybricare 46X Media (46X) from Sigma-Aldrich, USA; Neratinib (Cat #A8322) from APExBIO, USA . Any complete media (DMEM-HG or 46X) is formulated from the basal media by adding 10% FBS and 1% P/S.

### μFTA and Valve - Design and Manufacture

The μFTA was designed first by drawing the masks using Adobe Illustrator and KLayout. The 3-layer µFTA device comprises three separate layers - the top feeding, bottom tissue-seeding, and middle porous membranes as the integral parts. Trapping mechanisms (“catchers”) are integrated into the tissue-seeding layer to trap the tissue pieces within individual chambers. Various designs were created and tested before settling on a final prototype, in formats of 4x2, 4x4, or 8x4 arrays. The devices were manufactured through standard soft photolithography microfabrication techniques at our core fabrication facility (Nanofabrication Facility at the Advanced Science Research Center, CUNY, NY, USA), using negative photoresist SU-8 (Microchem, USA) as the molding photoresist and polydimethyl-siloxane (PDMS, Dow Corning, USA) as the casting material.

We used PE/3 tubing (Scientific Commodities, Inc, USA) for media flow, and 3/8in x 5/8in (ID x OD) medical grade Tygon tubing (US Plastics Corp., USA) for tissue seeding. The manufactured devices were tested with food dye or isopropyl alcohol (IPA) for leaks and to ensure fluid perfusion across the middle interface layer. The devices were then fully flushed with ethanol (70%v/v) and exposed to UV light for 1 hour for sterilization.

A physical valve was implemented into a cylindrical section cut away from a fully fabricated 4x2 array format μFTA (Fig. 1.G(i)). The cut-away section is located along the channel in between two rows on the bottom layer and was not removed from the μFTA. A leak-proof adapter was built atop this cut-away section, and a hermetic seal below it. The top adapter is structured to allow a plug (push button) to be pushed through it, dislocating the cut-away section of the device, by obstructing the bottom channel. The hermetic seal under the cut-away is a 0.5-1mm deep PDMS well that is structured within it, with a thin piece of glass (No. 2, Corning) bonded underneath for rigidity. The bottom seal and top adapter were then bonded under and over the valve cut-away section, respectively, using PDMS as an adhesive. Leakage was quantified by flowing two solutions at 1ml/day: (i) 100%: 10µg/ml Doxorubicin in 1X Phosphate Buffered Saline and (ii) 0%: just PBS. The two solutions are flowed parallel to each other across the valve, and the fluorescence of the doxorubicin can be measured using 470/560 nm excitation/emission spectra to quantify dox levels leaking across the valve. The flow is maintained for 24 hours. *As per the schematic shown in Fig. 1.D.(v), the 100% input chamber C4 and 0% input chamber C5 for the non-valved system are monitored for cross-talk of doxorubicin, representing the corresponding chambers that would be immediately across the valve. Thus, chambers C9 and C10 for the valved system are monitored for any doxorubicin leakage*.

### CFD optimization

#### Computational Fluid Dynamics Model

The CFD software used for the simulation was COMSOL v5.0.The simulation was set up with standard physical parameters and a previously measured glucose consumption rate [45]. Briefly, standard water density (∼1g/ml), standard diffusion coefficients for glucose, O_2_, and CO_2_ in water and PDMS were used (on the order of 10^-9^ m^2^/s). The glucose consumption rate as estimated from a previously conducted cell culture experiment is 5.2721x10^-5^ mol/(m^3^*s). The device was modeled using three feeding rates: receiving 1ml/day, 5ml/day, and 10ml/day of the appropriate media (glucose concentration of 1g/L). A central, elliptical cylinder (approximate volume of 263μl) was used as a region to model the tissue sample with a permeability of 10^-15^m^2^. The laminar flow module (Brinkman model) coupled with a Dilute Species Reaction module in COMSOL were used.

### Flow Verification via Particle Imaging Velocimetry (PIV)

Flow profile in the device was recorded through fluorescent particle tracking (particle imaging velocimetry, or PIV). Orange fluorescent latex beads of 2μm mean diameter and 1.012g/cm^3^ mean density (Sigma-Aldrich, USA) were suspended in water and flowed into the μFTA at various velocities corresponding to the 1,5,or 10 ml/day CFD simulations. The epifluorescent particles were tracked by our AxioObserver Z.1 microscope (Carl Zeiss), using time lapse or fixed framerate videography. The particles are traced over a certain time frame (4,10, or 50 seconds). Traces were tracked in Blender v2.7.1 or Adobe Photoshop CS6 (Adobe, Inc.), segmented in MATLAB v7 (Mathworks, Inc.), and matched to the framerate to calculate particle velocity. This velocimetry data was compared to CFD simulation *under free flow* conditions (without any tumor piece) to validate simulated flow profiles against observed recordings.

### Cell-line Derived Tumor Xenograft Models

#### Cell-derived (CDX) Models for device validation

MDA-MB231 triple negative (ER-/PR-/HER-) cancer cells (ATCC, USA) were cultured in complete high-glucose Dulbecco’s Modified Eagle Medium (DMEM-HG), formulated with 10% FBS and 1% Penn-Strep. MB231 cells with Green Fluorescent protein (GFP) were obtained from the Jiang Lab at MSKCC. Cells were inoculated by injecting 5 x 10^6^ cells in 50% matrigel subcutaneously in the left and right flanks of 4-6-week-old female NOD/SCID mice. Tumor growth was externally monitored daily via calipers. Mice were sacrificed when the tumor exceeded a 10mm diameter and tumors were harvested for ex vivo μFTA device experiments. BT474 (HER2+) cells with or without Red Fluorescent Protein (RFP) expression were similarly used to generate the HER2+ CDX model.

### Patient-Derived Tumor Xenograft Models

#### Patient derived (PDX) Models

M271(ER+/ PR- /HER2+), M37 (HER2+), and M156 (triple negative breast cancer, TNBC) are xenograft models that were derived from patient biopsies clinically characterized as breast cancer at the Chandarlapaty lab at MSKCC. The generation and propagation of M37 was described in a prior work[46]. M271 and M156 PDXs were obtained from patients who sign an informed consent form according to the Human Biological Samples Policy. Six to eight week-old female NOD *scid* gamma (NSG) mice [47] were purchased from the Jackson Laboratory . Three days before tumor implantation took place, estrogen pellets (0.18mg) were implanted subcutaneously into one flank of each mouse. Patient biopsies were dissected into thin 3x5 mm pieces and subcutaneously implanted into the opposite flank with a 10-gauge trochar. Animals were monitored daily for visible tumor growth, animal weights and tumor measurements were performed weekly to monitor changes in health. After the tumors reached 150 to 200 mm^3^, the mice were sacrificed, and the tumors were excised in a sterile manner. Tumor re-implantation was performed similarly. This PDX establishment protocol was approved by the IACUC review board at MSKCC.

### Tissue transport, sizing and seeding into the µFTA

All harvested xenograft tumors (PDX or CDX) were transported in ice-cold PBS or RPMI media (with 1% Penn-Strep) within 45 minutes from MSKCC to CCNY. Each explanted tumor, about 0.5-1 cm diameter ellipsoids, were placed into a 10-cm petri dish with a 3-5ml mix of complete DMEM medium and PBS in a 1:1 ratio (DMEM-PBS) at room temperature. Each explant sample was carefully minced manually to around 0.5mmx0.5mmx0.5mm sized micro-dissected tissue pieces. These microtissues were seeded into the µFTA via syringe as a suspension of 20-30 pieces per 5ml of media.

The sterilized µFTA device was primed with IPA, then flushed with sterile DD water, PBS, and complete or serum-free media sequentially. A syringe containing 20-30 pieces of microtissues suspended in ∼5ml of media was dispensed into the bottom layer at a controlled rate of 50-200 µl/sec. The pieces are secured in place by the catchers within the device µFTA. After tissue seeding, fresh complete media was introduced into the µFTA through the top and bottom layers. The outlets were sealed with waterproof inserts constructed of PDMS-filled tubing. The µFTA was then ready for culture via continuous media supply through the feeding layer.

### Tissue Culture and Treatment

Our experimental conditions have been briefed in the following table (all conditions implicitly have carrier controls, and were cultured in the μFTA unless specified):

**Table 1:** Culture conditions for each of the cell-derived xenograft (CDX) or patient derived xenograft (PDX) samples. #N = Number of days in normal culture media; #D = number of days in drug containing media; Ner = Neratinib, Dox = Doxorubicin. TC = Tissue Culture. MDA-MB 231 (CDX) and M156 are triple negative(ER-/PR-/HER2-). BT 474 (CDX), M37 (PDX), and M271 (PDX) are HER2+, but only M37 is Neratinib resistant. “With GFP/RFP” indicates cells were transfected with GFP or RFP expressing genes.

| ***Xenograft Type*** | ***HER2 status*** | ***Culture Condition (#N or #D)*** | ***Culture Substrate*** | ***Assays/Stains*** |
| --- | --- | --- | --- | --- |
| **<u>MDA-MB 231</u>**<br>- GFP | - | 7N+4D Dox | μFTA | Micrograph Observation |
| MDA-MB 231 | - | 3N+4D Dox (2μg/ml, 20μg/ml)<br>3N+4D Ner (180nM, 1800nM) | μFTA | Viability |
| **<u>BT-474</u>** - RFP | + | 7N | μFTA | Hoechst dye |
| BT-474 | + | 3N+4D Ner (180nM, 1800nM) | μFTA | Viability |
| **<u>M156</u>**<br>(BRCA1 mutant) | - | 7N | μFTA | Viability, Proliferation |
| **<u>M37</u>**<br>(Ner resistant) | + | 4N+3D Dox (2μg/ml, 20μg/ml) | TC well plate, PDMS-coated surface, μFTA | Viability, Proliferation |
| **<u>M271</u>**<br>(Ner sensitive) | + | 3N+4D | μFTA | Viability, Proliferation, Histology |
| M271<br>(Ner sensitive) | + | 7 days after tumor injection, 3 days/week p.o. of Ner (20mg/kg), up to 36 days | In-vivo (mice) | Physical measurement, Western Blot |

### Functional Assays

All functional assay staining dyes were obtained from Fisher Scientific, USA, branded as Invitrogen™ Molecular Probes™, unless otherwise stated. The live/dead dyes used were Calcein-AM, Ethidium Homodimer-1, Hoechst (FluoroPure™ Hoechst 33342, Trihydrochloride, Trihydrate), while proliferation was assayed using the Invitrogen™ Molecular Probes™ Click-iT™ Plus EdU Alexa Fluor™ 555 or Alexa Fluor™ 488 Imaging Kit. Stock solutions for Calcein-AM and Ethidium Homodimer (EthD-1) were prepared as 2µg/µl formulations in DMSO. Hoechst 33342 was available as an aqueous stock solution of 1 mg/ml (16mM). For the µFTA, we use a working concentration of ∼ 1:250 dilution each of Calcein-AM and EthD-1, and ∼1:500 dilution of Hoechst in prepared in complete (CalEthHt-CM) or serum-free media (CalEthHt-SF) in the appropriate tissue culture media (DMEM-HG or 46X).

For viability assays, the µFTA is flushed through the feeding layer with 2.5-3ml of CalEthHt-CM prepared above at 200-300µl/min for the first 1.5ml, and then at 1ml/hr for 40 minutes. The dispensing solution is then changed to CalEthHt-SF and perfused for 1 hour at 1ml/hr. The µFTA is kept in the incubator at 37°C and 5% CO_2_ culture conditions during the staining steps. Finally, the µFTA was perfused with PBS for 1.5-2 hours at 1-1.5ml/hr to clear the staining solution and media.

For proliferation assays, the final EdU staining cocktails are formulated as per manufacturer’s protocol in 2.5ml batches. Tissue samples were cultured 16-18 hours in EdU to obtain distinctly stained signals. The 1000X EdU stock was diluted to a 2X dilution in complete media (DMEM-HG or 46-X) and replaced the normal culture media for overnight perfusion. The media was cleared out by washing with PBS for 25-30 minutes@100µl/min, followed by 4% PFA fixing for 2 hours@50-75µl/min. Any intermediate washing or blocking steps recommended in the manufacturer’s protocol for adherent cells on a slide was adopted as a 75µl/min flow rate into the µFTA, for a minimum of 30-45 minutes. The 2.5ml EdU staining cocktail was perfused over a duration of 1.5-2 hours in the µFTA.

Samples for immunohistochemistry of Neratinib treated M271 cells were sent for processing to the MSKCC core facility, for standard hematoxylin and eosin (H&E) staining, along with TUNEL, Ki67, and HER2 immunostaining. The stained samples were subsequently analyzed via the expertise of two breast pathologists (FP and LCC).

*Static Culture/Out-of-Device Staining*. Staining on tissue culture well plates were done at a similar time scale and concentration as the µFTA. For viability staining, tissue samples were kept on an orbital shaker set to 60 rpm with CalEthHt-CM for about 1 hour, followed by a PBS wash for another hour on the shaker. For proliferation staining, tissue samples were first placed in well plates on an orbital shaker at 60 rpm with media containing 2X EdU for 2 hours in the incubator. The plates are then moved off the shaker to be left in the incubator overnight. The period of staining on the culture plate and µFTA was kept the same. The EdU staining cocktail was retained in the wells for 1.5-2 hours on the 60 rpm shaker.

*Image Analysis.* Tissue fluorescence image analysis algorithms were developed in-house in MATLAB v.2017b for automated segmentation and quantitative analysis, and an end-result graphic flowchart is depicted in Supp. Fig. 1. Optimized segmentation algorithms were used to differentiate live, active regions from dead regions. The tissue was also categorized into “peripheral” and “core” regions, where the peripheral region is defined up to a depth of ∼100 µm from the surface, and the “core” as deeper regions. A nonlinear weighting compensates for depth-dependent signals. For “whole tissue” measures, both the “core” and “periphery” were included.

*Statistical Analysis of Image Data.* Quantitative live/dead or proliferation signal data acquired through the steps in “Image Analysis” section above are analyzed using the non-parametric Kruskall-Wallis test, followed by a post-hoc Dunn-Bonferroni test using Microsoft Excel. Since our samples do not strictly follow a parametric profile, we choose the non-parametric analytics over standard ANOVA.

## Acknowledgements

Funding was provided by: (1) NIH/NCI U54 CA132378 (2) Pershing Square Sohn Prize for Young Investigators in Cancer Research, (3) NSF TI 1938430, and (4) CUNY ASRC CAT Program.

We would also like to acknowledge the substantial efforts and contributions of Dr. Helen Kang and Dr. Jiao Wu from the Jiang Lab at MSKCC. Dr. Helen Kang has been pivotal at the initial stages of the project when we tested our µFTA with CDX samples, which paved the way for great progress with PDX samples which are at the centerpiece of this paper. Dr. Jiao Wu has offered her technical expertise in preparing and analyzing histology sections and providing more CDX samples.

## Notes

### Competing Interest Statement

The authors have declared no competing interest.

### Summary of Updates

Figures 3 and 4 were missing their significance bars, as the graphs without the significance marks were uploaded. The proper graphs with the significance bars have been uploaded.

