## Supplemental Figures for "Microfluidic Tissue Array Platform for Personalized Drug Screening Using Tumor Explants or Biopsies"

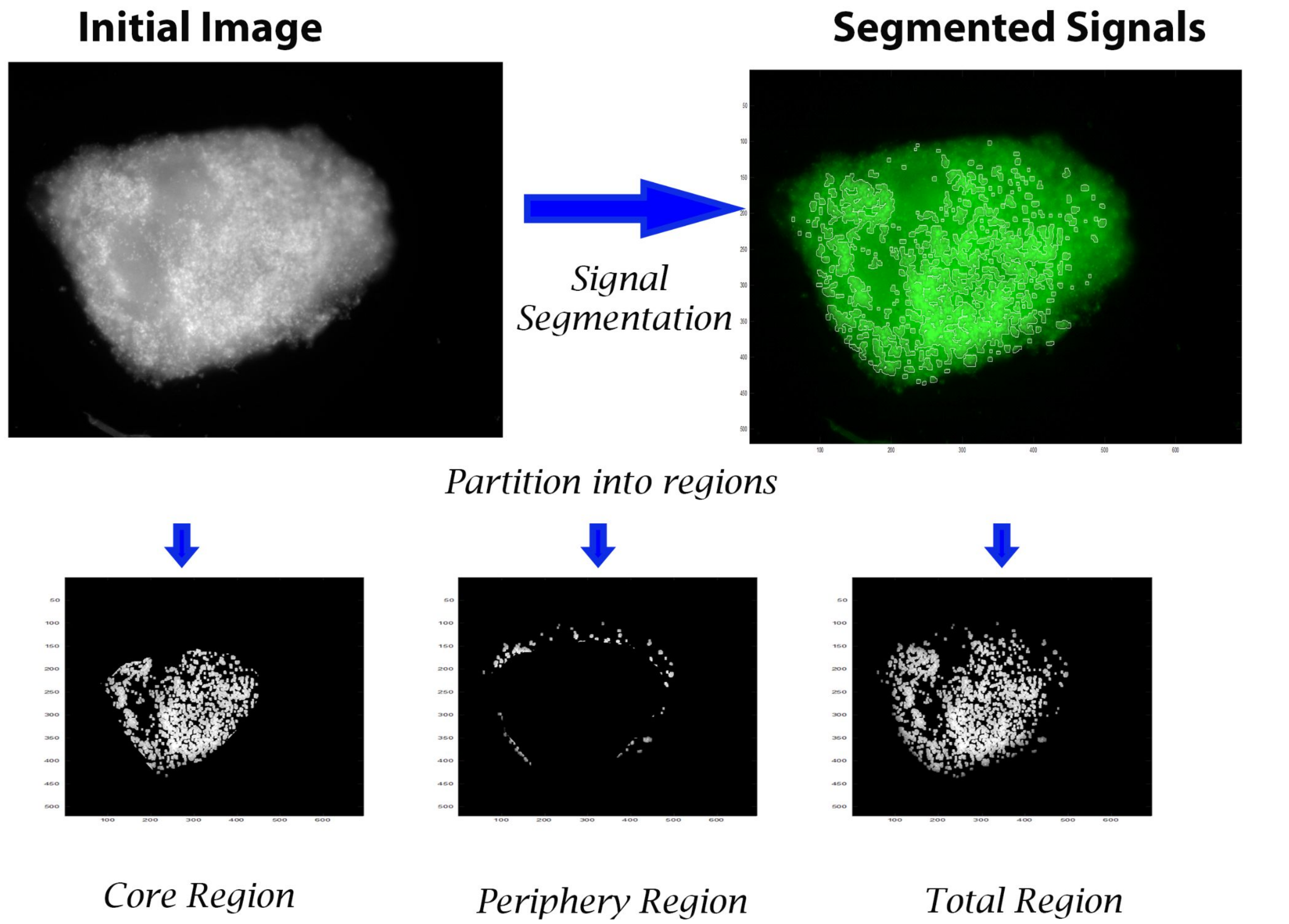


### Supplementary Figure 1

**Supp. Figure 1: Demonstration of signal extraction from fluorescent tissue images.** Intricate spatial filters are used to segment the cellular signals within the tissue (top left), with the segmentation shown as small bounding regions within the tissue. The image is pseudo-colored green from the grayscale for better visibility. The identified tissue is then partitioned into a peripheral (~100µm deep) region and the core region. The final segmented pixels of the core/peripheral cell signals are indicated in a black/white mask. The “total region” is the union of the core and peripheral masks.

### Supplementary Figure 2

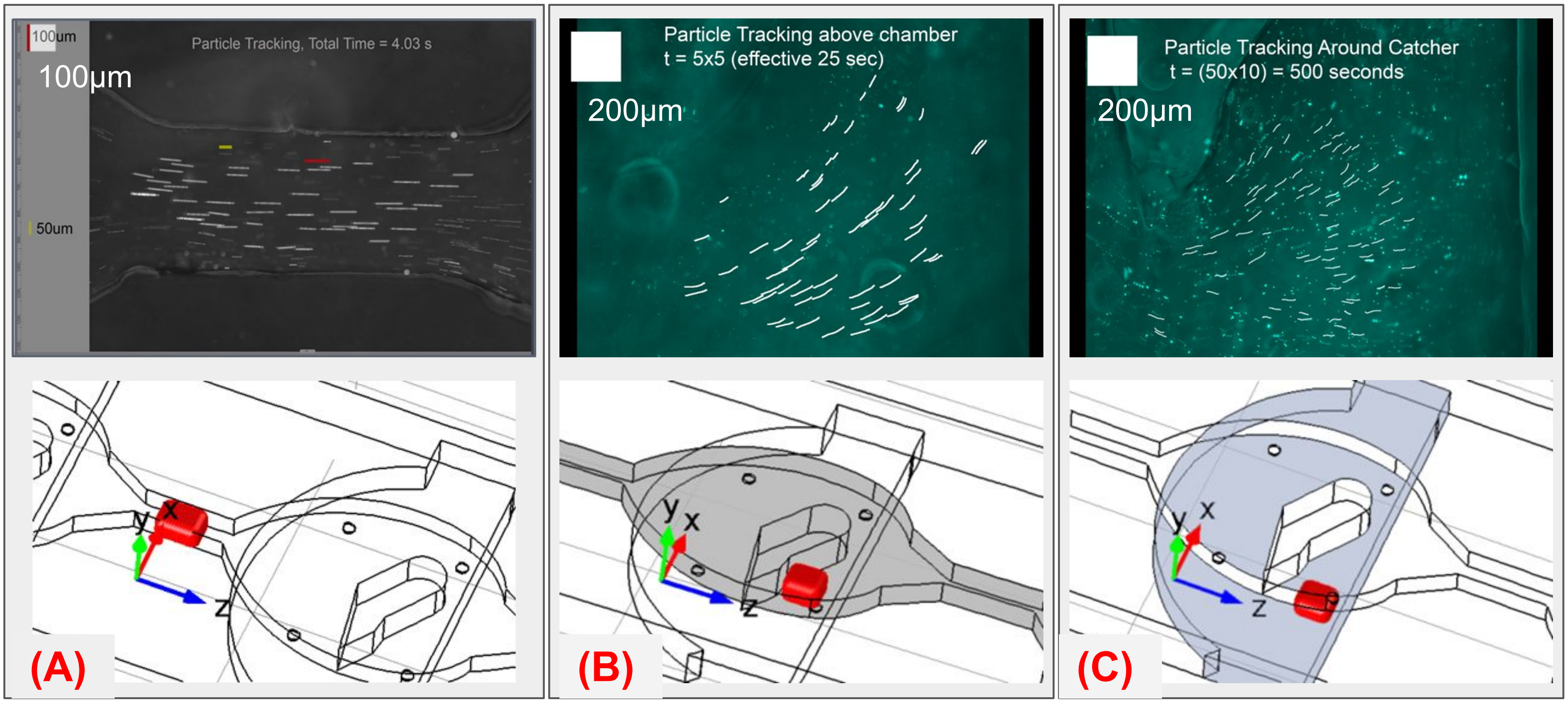


| Location | (A) Top Channel (between chambers) | (B) Top Channel (above chamber) | (C) Bottom Layer (around chamber/hole) |
|---|---|---|---|
| **Speed (From COMSOL)** | 21.7(±3.38) μm/sec | 2.83(±0.76) μm/sec | 0.103(±0.019) μm/sec |
| **Speed (from particle flow)** | 22.7(±3.35) μm/sec | 2.73(±0.32) μm/sec | 0.113(±0.067) μm/sec |

**Supp. Figure 2: Particle Velocimetry and Velocity Profile from Computational Fluid Dynamics Simulation (CFD).** The top row images in (A), (B), and (C) show the locations in the μFTA where 2μm fluorescent latex beads were used to measure velocity within the device at the locations identified in the bottom row on the 3D CAD drawing. (A) is the region which only has the top channel, and is the media feeding path that connects between chambers. (B) shows the region in the top chamber above where the tissue sample would sit in the μFTA during culture. (C) Shows the region within the catcher where the tissue sample would actually be seated during the culture duration in the bottom layer. The table below shows the summary of the average velocity of the particles in velocimetry compared to the expectation from the same probing regions shown in (A),(B), and (C).

Supplementary Figure 3

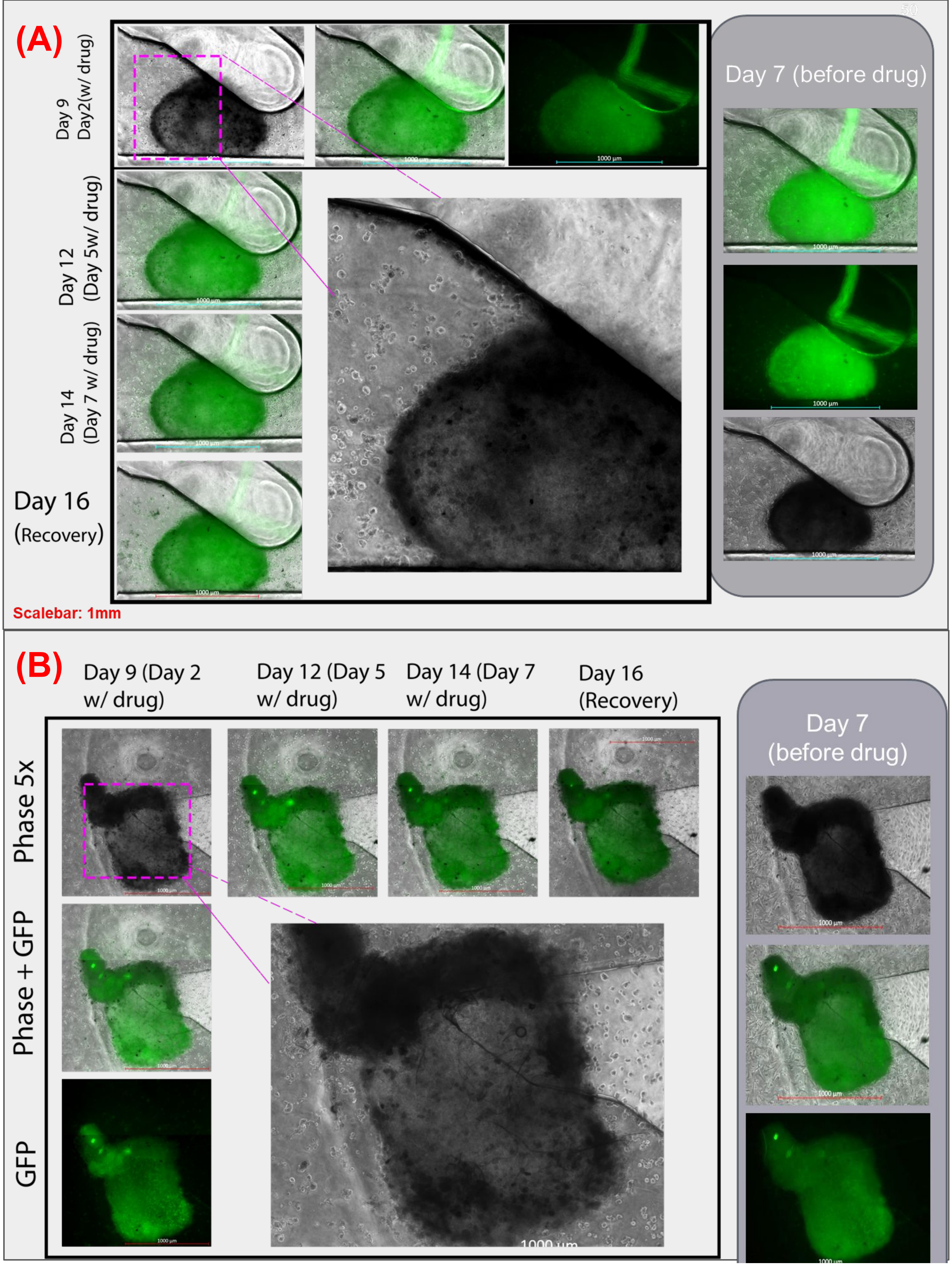


**Supp. Figure 3: Tissue Clearance on drug treatment after 7 day culture.** Two samples of triple-negative MB-MDA 231-GFP xenograft biopsies microdissected and cultured on the µFTA for 7 days (right column in A and B), after which doxorubicin treatment is induced in the flowing media for 7 days, with images showing days 2, 5 and 7 of drug treatment. The tissue is cleared out of cells by day 2, as shown in the expanded insets for (A) and (B), and fluorescence photomicrograph 2 days after the drug is stopped (Day 16) is also shown.

### Supplementary Figure 4

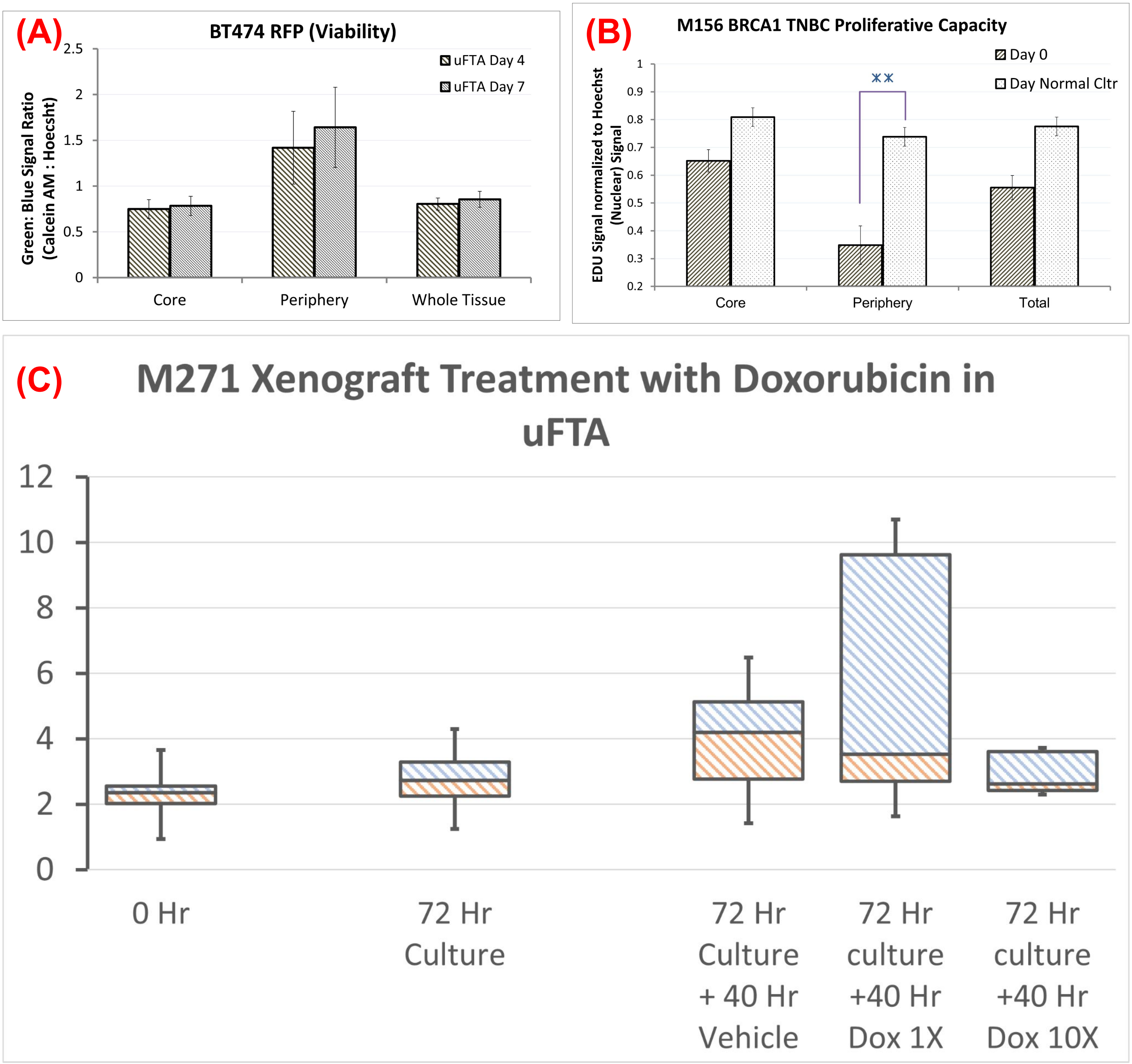


**Supp. Figure 4: Tissue Growth and Drug testing on the μFTA - other tissues/preliminary runs.** (A) BT 474 xenograft tissues were cultured in the μFTA as a representative cell line for HER2+ breast cancer and probed with a viability assay. (B) M156 Triple negative breast cancer (TNBC) was a patient-derived xenograft (PDX) used as a preliminary run for TNBC from a patient, instead of a cell line and then probed with a proliferation assay at the end of 7-day culture. (C) M271 is a HER2+ PDX, which is not responsive to doxorubicin, as indicated by a 3-day culture (72 hours) followed by by ~2 days (40 hours) of 1X and 10X doxorubicin doses.

Supplementary Figure 5

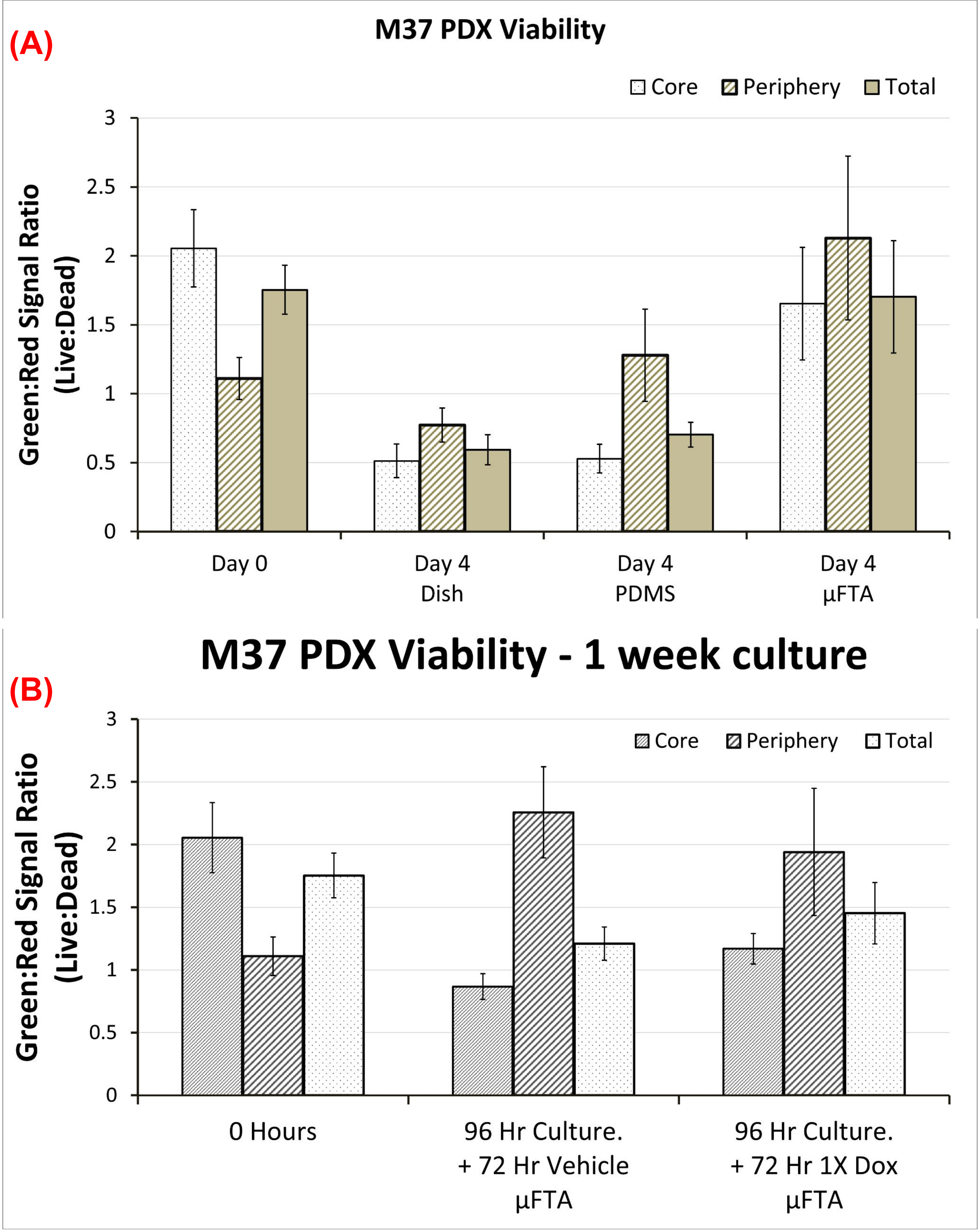


**Supp. Figure 5: M37 (HER2+, neratinib sensitive) tissue growth and drug testing on the µFTA and/or other standard culture platforms.** (A) Shows the core, peripheral and total viability of M37 tissue on a standard culture well plate, a well plate coated with PDMS, and the µFTA, comparing across 4 days of culture to Day 0. (B) Shows doxorubicin treatment effect compared to no drug ("vehicle") on the viability of M37 tissue after 4 days of culture (96 hours) followed by 3 days of treatment (with or without drug, 72 hours).

### Supplementary Figure 6

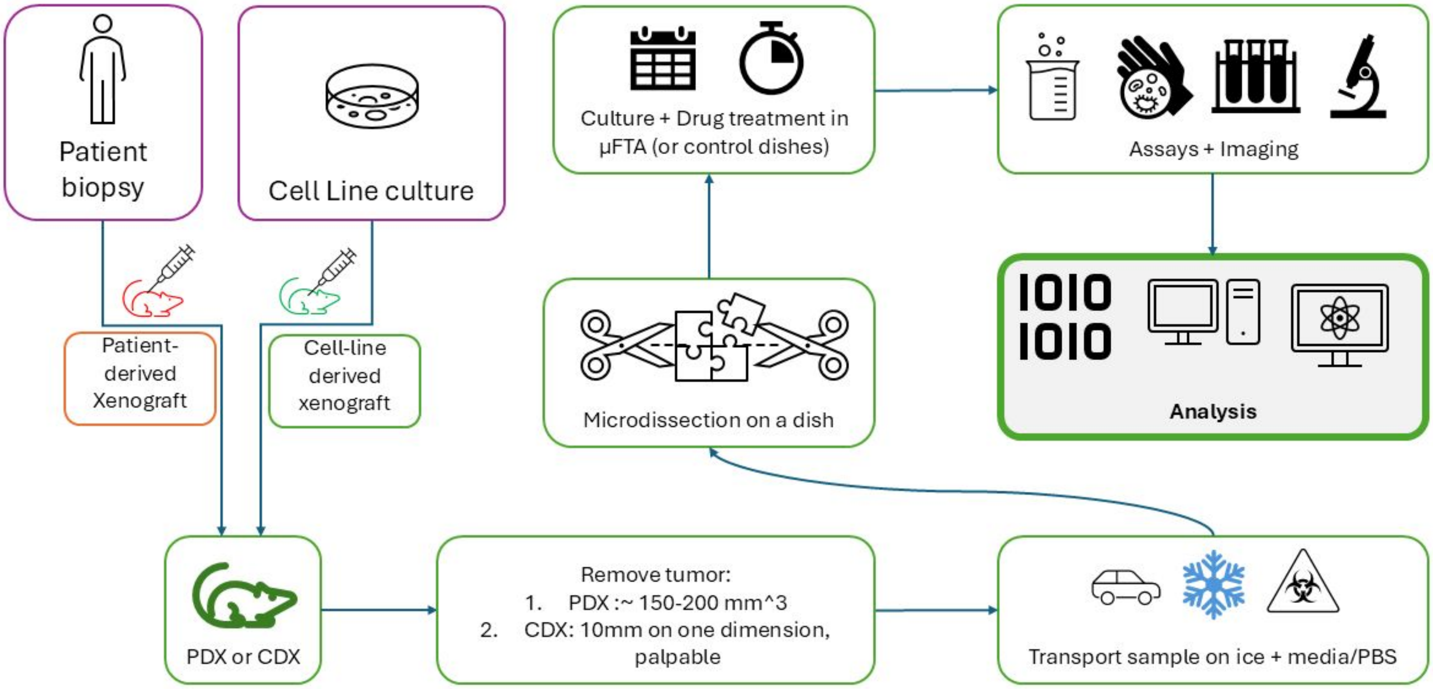


Some images will be replaced with custom images to avoid any copyright. Icons from the Microsoft Suite seem to be copyright-free.
